# Automated hierarchical block decomposition of biochemical networks

**DOI:** 10.1101/2025.03.07.642055

**Authors:** Manvel Gasparyan, Satya Tamby, G.V. HarshaRani, Upinder S. Bhalla, Ovidiu Radulescu

## Abstract

Biochemical networks are models of biological functions and processes in biomedicine. Hierarchical decomposition simplifies complex biochemical networks by partitioning them into smaller blocks (modules), facilitating computationally intensive analyses and providing deeper insights into cellular processes and regulatory mechanisms. We introduce a novel algorithm for the hierarchical decomposition of large-scale biochemical systems. By using causality and information flow as organizing principles, our approach combines strongly connected components with *r*-causality to identify and structure manageable network blocks. Benchmarking against a comprehensive database of biochemical reaction networks demonstrates the computational efficiency and scalability of our algorithm. To ensure broad applicability, we integrate our algorithm into tools that support standardized Systems Biology Markup Language (SBML) formats, facilitating its use in biochemical modeling workflows.

## 1 Introduction

In network analysis, understanding hierarchical structures within directed networks is crucial for deciphering organizational dynamics and information flow in these systems.

Chemical Reaction Networks (CRNs) are used to model the functions of biological cells in fundamental biology and health-related applications. CRNs take the form of oriented bipartite graphs, with one set of nodes representing molecular species and another set representing chemical reactions.

To simplify the comprehension of CRNs, biologists often break down large networks into more manageable pathways. These sub-models, commonly referred to as blocks or modules, are assumed to exhibit simpler functions. There are currently several public biological networks repositories databases in which the cellular wiring is presented as functional modules (pathways) [1–3]. However, decomposition into functional modules is typically performed manually and lacks clear mathematical principles.

Computational analysis of these complex models involves challenging problems in machine learning [4, 5], constraint programming [6], and formal verification [7, 8], all of which could benefit from network decomposition. Network decomposition also helps understanding dynamical properties of networks, such as multi-stationarity and robustness [9], stability [10–12], species persistence [13].

Significant efforts have been made to formalize the decomposition of CRNs and other biological networks. One popular approach is based on community detection, which identifies sets of nodes that interact more frequently with each other than with the rest of the network [14, 15]. Strongly Connected Components (SCCs) represent a specific type of community, where each node has a direct or indirect interaction with every other node in the group [16, 17].

While community detection highlights local network features, hierarchical decomposition emphasizes the global organization of the network [18]. Hierarchies are commonly observed in social networks, where certain individuals act as leaders and exercise control over others [19]. Similar organizational patterns can be found in biological systems [20]. For instance, signal processing within cell molecular pathways or neural networks in neuroscience often follows hierarchical principles, where molecules or neurons serve as receptors that transmit signals through various processing layers to effectors [5, 21, 22].

In graphical models, hierarchies are constructed with causality in mind: nodes with lower rank control those with higher rank. Identifying a hierarchy involves ranking the nodes of oriented graphs, such that the sources have the lowest hierarchical rank, and edges go from lower to higher rank nodes. Such ranking is possible for Directed Acyclic Graphs (DAGs). However, real networks seldom conform to the structure of DAGs. For networks that contain cycles, hierarchies can be identified by minimizing agony, a measure that increases with the number of edges for which the source node has a higher hierarchical rank than the target node [19,23]. Identifying the hierarchical organization of systems is particularly useful in the growing field of causal machine learning [24].

Because agony based methods rely on optimization, their output suffers from degeneracy and randomness, especially when the network has multiple cycles. Furthermore, it may be very different from the biological intuition. Finally, biological networks have multi-scale organization, with hierarchies occurring at various scales (meaning reactions, sets of reactions, pathways, see [21, 22]). In order to increase the robustness of the decomposition and bring them closer to biologist’s intuition we propose hierarchical decomposition methods of CRNs that combine strongly connected components, agony and the notion of *r*-causality that handles scale. The *r*-causality means disregarding influences that span more than *r* steps in the interaction graph, a principle previously applied to community detection [25].

In this paper, we introduce an algorithm for the hierarchical block decomposition of CRNs. The algorithm begins by constructing the interaction graph of the CRN, a directed graph representing the interactions between species participating in reactions. Depending on a pre-specified parameter *r*, the algorithm proceeds in one of two directions. When *r* is infinite, it employs a strict definition of causality and computes blocks as SCCs. For finite values of *r*, blocks are *r*-Strongly Connected Components (*r*-SCCs), a concept derived from SCC in which interaction propagation is restricted to paths of length at most *r*. The blocks are used to define a hierarchical block decomposition of the network. The algorithm is implemented as a Python tool called **BlockBioNet**. We further demonstrate how our decomposition facilitates the generation of an informative and aesthetically structured visualization of CRNs. Specifically, we define a layout that enables three distinct representations: (i) the species-reaction graph, illustrating how species participate in reactions; (ii) the interaction graph; and (iii) the quotient graph of the interaction graph defined by our block decomposition. The CRNs visualization is implemented as a Python tool called **CRNPlot**.

## 2 Preliminaries

We present the preliminary notions necessary for our automated hierarchical block decomposition method, beginning with a standard framework for modeling CRNs, followed by a hierarchy detection method developed for social directed graphs in [19]. Although originally designed for social graphs, this method can be adapted to any directed graph, including the ones representing CRNs.

### 2.1 Chemical reaction networks as dynamical systems

A CRN involving *s* distinct chemical species, denoted as 𝒳 _*i*_, *i* = 1, …, *s*, and *n* distinct (bio)chemical reactions, denoted as ℛ_*j*_, *j* = 1, …, *n*, can generally be represented as:

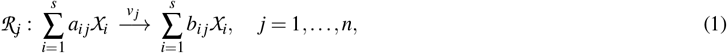

The net changes in the concentration of species, form the elements of the stoichiometric matrix. For a chemical reaction network described by Eq. (1), the stoichiometric matrix **S** is an *s* × *n* matrix where *S*_*i j*_ = *b*_*i j*_ − *a*_*i j*_. Positive entries *S*_*i j*_ *>* 0 indicate the production of the *i*^th^ species in the *j*^th^ reaction, while negative entries indicate its consumption.

Each reaction is also characterized by its rate, that is a function of the concentrations of the species, *v*_*i*_: ℝ^*s*^ → ℝ. This type of function is called a rate law and can take various forms, such as mass-action, Michaelis-Menten, or Hill, to name just a few examples.

For the *j*-th reaction, species such that *S*_*i j*_ *<* 0 and *S*_*i j*_ *>* 0 are called *reactants* and *products*, respectively. Any *i*-th species that is neither reaction nor product (i.e. *S*_*i j*_ = 0) but whose concentration influence the reaction rate (i.e. 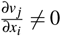) is called *modifier*.

Species concentrations evolve in time as a result of chemical reactions. In chemical kinetics, under the well-stirred reactor assumption, they satisfy the following system of Ordinary Differential Equations (ODEs):

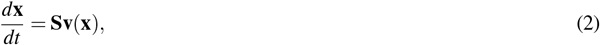

where 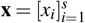 is the vector of concentrations and 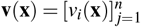 is the vector of reaction rates, which is a vector-valued function of the species concentration vector **x**, that is, **v**: ℝ^*s*^ → ℝ^*n*^.

#### Example 1.

*Consider the following example of a CRN consisting of five distinct chemical species, denoted as* 𝒳_*i*_, *i* = 1, …, 5, *and three unidirectional reactions given by:*

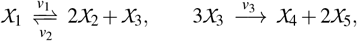

*Under the mass-action kinetics assumption, the reaction rates are:*

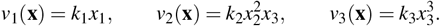

*The stoichiometric matrix* **S** *is:*

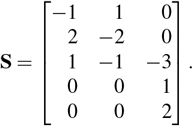

*Accordingly, the system of ODEs governing the dynamics of the species concentrations can be formulated as follows:*

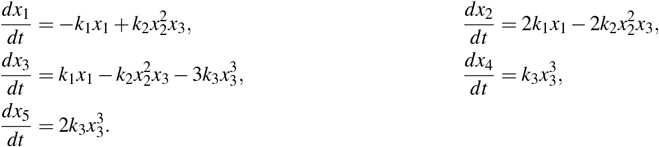

Other examples can be extracted from **ODEBase**, an extensive database of ODE representations of CRNs, formatted for symbolic computation [26].

### 2.2 Agony-based hierarchy detection in directed graphs

Here we outline a method for identifying hierarchies in directed graphs with cycles, as detailed in [19].

In the context of network analysis, we examine a directed graph 𝒢 = (𝒱, ℰ), where 𝒱 denotes a finite set of nodes and ℰ denotes a finite set of directed edges, potentially forming cycles. Each node *v* ∈ 𝒱 is assigned an integer rank ω(*v*) by a rank function ω: 𝒱 → ℕ. Nodes can share the same rank. These ranks establish a hierarchical relationship: nodes with lower ranks are expected to influence or direct nodes with higher ranks, resulting in directed edges *u* → *v* when ω(*u*) *<* ω(*v*). Conversely, edges *u*→ *v* are less prevalent and merit attention when ω(*u*) ≥ ω(*v*), reflecting a less optimal direction of influence based on the assigned ranks.

These requirements are formalized as an optimization problem. The agony in the graph 𝒢 relative to the ranking function ω is defined by:

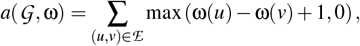

and the minimum agony in the graph 𝒢 is the least agony achievable over all possible rankings, i.e., *a*(𝒢) = min_ω_ *a*(𝒢, ω). The hierarchy in graph 𝒢 is defined as:

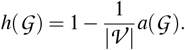

Given that for any ranking function ω, the number of edges in the graph 𝒢 serves as an upper bound for the agony in 𝒢 relative to ω, the hierarchy falls within the interval [0, 1].

To find the hierarchy for any directed graph, we need to search over all rankings and find one that gives the highest hierarchy. In [19], the authors provided an efficient *O*(*nm*^2^) algorithm of finding this ranking function for any graph with *n* number of vertices and *m* the number of edges. A more efficient *O*(*m*^2^) algorithm was proposed in [23]. In Figure 1, directed graphs are presented with nodes ranked according to this method. In graphs where a perfect hierarchy is not achieved, edges that violate this hierarchy are highlighted in red to indicate discrepancies.

**Figure 1:**
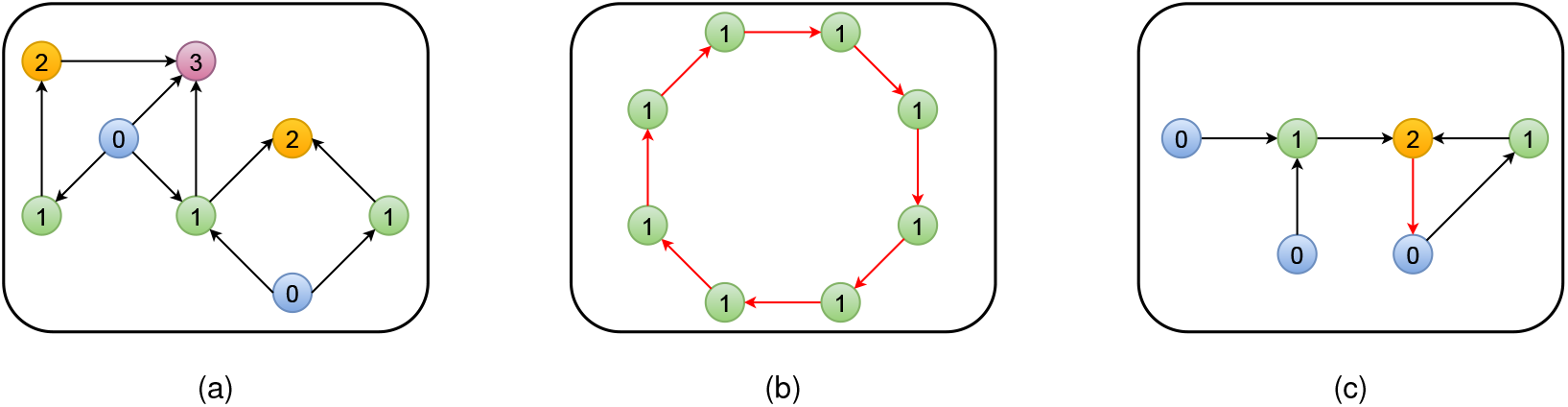
Examples of directed graphs with their hierarchies. The edges violating the hierarchies are highlighted in red. (a) Acyclic graph: *h*(𝒢) = 1. (b) Simple cycle: *h*(𝒢) = 0. (c) Cyclic graph: *h*(𝒢) = 1*/*2.

## 3 Hierarchical block decomposition

This section introduces the key concepts of Autonomous Nested Hierarchical Decomposition (a-NHD), and *r*-Causal Nested Hierarchical Decomposition (r-NHD), two related hierarchical block decompositions.

### 3.1 Nested hierarchical decomposition

First, let us introduce the concept of an autonomous pair. These are pairs of subsets of species and reactions, defined based on their intrinsic causal relationships, and they do not require additional elements to compute their time evolution.

#### Definition 1

(Autonomous pairs). *An* autonomous pair *is a subsets pair* (*I*, *J*) ⊆𝒳 *×*ℛ *satisfying the following properties:*

1. *if a species is in the subset I*, *then all the reactions for which the net effect (production minus consumption) on this species is non zero are included in the subset* 𝒥. *In other words, for every* 𝒳_*i*_ ∈ *I*, *if S*_*i j*_ 0, *then* ℛ_*j*_ ∈ 𝒥.*1. If the model contains buffered species, i*.*e. species whose concentrations are kept constant irrespectively on the existence or not of chemical reactions acting on them, then they can be excluded from this condition*.
2. *if a reaction is in the subset J*, *then all the species on which the rate of this reaction depends are in the subset I*. *In other words, for every* 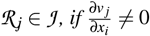, *then* 𝒳_*i*_ ∈ *I*.

The following theorem directly follows from the above definition.

#### Proposition 1.

*The indexed subvector* ***x***_*I*_ = [*x*_*i*_]_*i*∈*I*_ *satisfies the autonomous system of ODEs:*

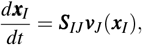

#### Algorithm 1

BlockBioNet decomposition.

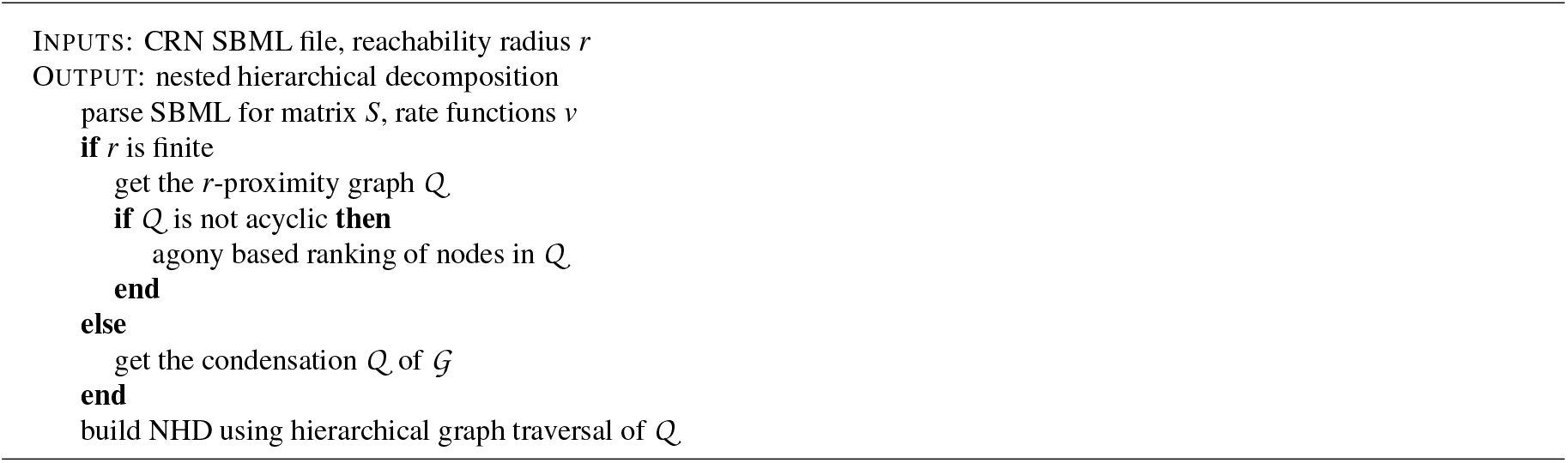

*where the matrix* ***S***_*IJ*_ *is extracted from the stoichiometric matrix* ***S*** *by selecting the rows I and the columns J*.

Proposition 1 implies that the concentrations of species in an autonomous pair can be determined from their initial values using only the reactions within the same pair. Naturally, the set of all species and the set of all reactions form an autonomous pair, but CRNs also contain smaller autonomous pairs. In applications such as hierarchical optimization [5], organizing autonomous pairs in a nested manner is beneficial. This leads to the concept of a-NHD.

#### Definition 2

(Autonomous nested hierarchical decomposition). *An autonomous nested hierarchical decomposition is a family of autonomous pairs* {(*I*_*m*_, 𝒥_*m*_): *m* = 0, …, *d* − 1} *satisfying the following properties:*

1. *I*_*m*_ ⊂ *I*_*m*+1_, *m* = 0, …, *d −* 1, *and I*_*d−*1_ = 𝒳.
2. 𝒥 _*m*_ *⊆* 𝒥 _*m*+1_, *m* = 0, …, *d −* 1, *and* 𝒥 _*d−*1_ = ℛ.

### 3.2 *r*-causality and *r*-strongly connected components

Causality is a key concept in the hierarchical decomposition of networks. In this section, we introduce the concept of species causality in CRNs through their interaction graphs, which illustrate the interconnections and dependencies between species. This form of causality will later be used to decompose a given CRN into blocks (modules), which will then be employed to determine the nested hierarchical decomposition of the CRNs.

#### Definition 3

(Interaction graph). *The interaction graph is a directed graph* 𝒢 = (𝒱, ℰ) *where* 𝒱 *is a finite set of nodes representing the chemical species, and* ℰ *is a finite set of edges defined by the following rules: if there is a reaction* ℛ_*k*_ *that consumes or produces the species* 𝒳_*j*_ *and the rate of this reaction is affected by the concentration of the species* 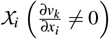, *then there is a directed edge in the graph from the species* 𝒳_*i*_ *(source node) to the species* 𝒳_*j*_ *(target node)*.

To compute the interaction graph, one needs the stoichiometric matrix and, for each reaction, the set of chemical species that influence the reaction rate. Because this information is time-independent, the CRN interaction graphs do not change over time, although weighted interaction graphs, that take into account interaction strengths, can be time dependent [27]. The interaction graph is used to define causality relations between species as follows.

#### Definition 4

(*r*-causality). *Let r be a positive integer, called the reachability radius. The species* 𝒳_*i*_ *is r-causal to the species* 𝒳_*j*_ *if there is a path of length at most r in the interaction graph* 𝒢 *from the species* 𝒳_*i*_ *to the species* 𝒳_*j*_. *We denote the r-causality of* 𝒳_*i*_ *to* 𝒳_*j*_ *by* 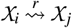. *We denote by* 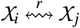 *the mutual r-casuality of* 𝒳_*i*_ *and* 𝒳_*j*_, *meaning that both* 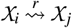 *and* 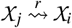 *hold*.

In this definition *r* plays the role of a scale parameter. The concept of *r*-causality refers to the idea that paths in the interaction graph exceeding a specified length *r* are ignored. This approach limits the scope of causality to short-range dependencies, which helps in focusing on the most relevant interactions within a given range. A similar notion, was used for community detection in [25] and named *p*-connectivity in that context.

Mutual *r*-causality is not inherently an equivalence relation. For example, if a species 𝒳_*i*_ is mutually *r*-causal with both 𝒳_*j*_ and 𝒳_*k*_, it does not necessarily imply that 𝒳_*j*_ and 𝒳_*k*_ are mutually *r*-causal to each other. To address this limitation, we introduce the concept of *r*-SCCs.

#### Definition 5

(*r*-strongly connected components). *An r-block is a maximal subset* ℬ ⊆ 𝒳 *in which every two species are mutually r-causal in the interaction graph* 𝒢. *An r-SCC is a lumping of r-blocks with non-empty intersections*.

Being in the same *r*-SCC establishes an equivalence relation among species, satisfying the criteria of reflexivity, symmetry, and transitivity. This framework offers a structured method to categorize and analyze species based on their multi-step interactions within the network, enhancing our understanding of the interconnectedness and dynamics of chemical species in complex systems. By identifying *r*-SCCs, we can accurately describe groups of species interconnected through multi-step interactions, providing a robust framework for analyzing the deeper structure of CRNs. This approach ensures that influences among species within each component are fully reciprocated up to the specified *r*-steps, reflecting a cohesive and interconnected subset of the network.

Determining the *r*-SCCs involves a process beyond straightforward partitioning. Initially, it entails computing the *r*-blocks and subsequently identifying those with non-empty intersections for lumping. To simplify this complex task, the mutual *r*-reachability graph provides an efficient alternative to the interaction graph.

#### Definition 6

(Mutual *r*-reachability graph). The mutual *r*-reachability graph ℋ *is an undirected graph in which the nodes represent species, identical to those in the interaction graph. The edges are defined as follows: an undirected edge exists between two nodes if the species corresponding to these nodes are mutually r-casual in the interaction graph* 𝒢.

If two species are connected by an undirected edge in the mutual *r*-reachability graph ℋ, this indicates that in the interaction graph 𝒢 there exist paths of length at most *r* connecting the two species in both directions. It follows from this definition that the set of species forming a connected component in ℋ also forms an *r*-SCC in graph 𝒢. Therefore, to determine the *r*-SCCs of 𝒢, we can identify the connected components of ℋ.

The set of *r*-SCCs forms a complete partition of the set of species, meaning they are disjoint, and their union is the entire set of species. Thus, this partition defines a quotient graph. In this quotient graph, each *r*-SCC corresponds to exactly one node. Additionally, there exists a directed edge from the *r*-SCC ℬ_*i*_ to the *r*-SCC ℬ_*j*_ if there is at least one edge in the interaction graph 𝒢 having a source node in ℬ_*i*_ and a target node in ℬ_*j*_.

#### Definition 7

(*r*-proximity graph). *The r-proximity graph* 𝒬 *is the quotient graph of the interaction graph* 𝒢 *defined by the set of r-SCCs*.

#### Remark 1

(*r*-causality for *r* → ∞). *It is important to note that we do not exclude scenarios where r becomes large, i*.*e*., *r* → ∞. *In this case, the concept of r-causality becomes strict causality. Two species are considered strictly causal if there exists a directed path from one species to the other, and vice versa. Consequently, in the limit r* → ∞, *the definition of the r-SCC becomes the same as the standard SCC definition. Furthermore, the r-proximity graph* 𝒬 *derived from* 𝒢, *based on the r-SCCs, effectively reduces to the condensation of* 𝒢 *–that is, the quotient graph of* 𝒢 *with respect to its SCCs*.

The *r*-SCCs (or SCCs in the *r* → ∞ case) correspond to blocks of species. The *r*-proximity graph (or the condensation of the interaction graph in the in the *r* → ∞ case) can be used to study the propagation of the interaction between blocks, emphasizing the presence of regulatory loops (feed-forward, feed-back) at the scale of the blocks.

The *r*-proximity graph 𝒬can be used to associate hierarchical ranks to all *r*-SCCs. If 𝒬 is acyclic, a rank of 0 is assigned to all sources (nodes with no incoming edge), while other nodes are assigned a rank equal to the maximal graph distance from the sources in 𝒬. If 𝒬 contains cycles, then hierarchical ranks are assigned to nodes using the agony-based method exposed in the Section 2.2.

The hierarchical ranks enable the definition of a r-NHD as follows:

#### Definition 8

(*r*-causal nested hierarchical decomposition). *A r-causal nested hierarchical decomposition is a family of pairs* {(*I*_*m*_, 𝒥_*m*_): *m* = 0,.. *where I*_*m*_ *is the union of all r-SCCs of rank smaller than or equal to m in* 𝒬, 𝒥_*m*_ *is the set of all reactions having nonzero net effect on species of I*_*m*_, *l* − 1 *is the maximal rank*.

From this definition it follows straightforwardly that

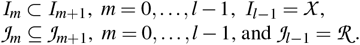

The two types of hierarchical block decompositions, the a-NHD and the r-NHD are related by the following proposition.

#### Proposition 2.

*When r* → ∞, *the sets* (*I*_*i*_, 𝒥_*i*_), *i* = 0, …, *l* − 1 *form a a-NHD in the sense of the Definition 8. This a-NHD is minimal in the following sense: if* 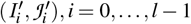 *is another a-NHD, then l* ≤ *d and* 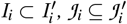 *for* 0 ≤ *i* ≤ *l* − 1.

In order to prove the Proposition we first prove the following

#### Lemma 1.

*For any SCCs component K of the interaction graph G and any subset I of an autonomous pair* (*I, J*), *one has either K* ⊆ *I or K* ∩*I* = ϕ.

*Proof of Lemma 1*. Consider that *K* ∩ *I* ≠ ϕ. Then, there is a species *x*_*i*_ that is both in *K* and *I*. According to the Definition 1, any species *x* _*j*_ such that (*j, i*) is an edge of 𝒢, should be in *I*. Furthermore, according to the Definition 1 and Remark1, any species strictly causal to *x*_*i*_, should be in *I*. But, all species in *K* are strictly causal to *x*_*i*_ by definition of SCCs, therefore, in this case, *K* ⊆ *I*.

*Proof of Proposition 2*. (*I*_0_, *J*_0_) where *I*_0_ is a source of the condensation of 𝒢 is an autonomous pair. Indeed, if a species *x*_*i*_ is in *I*_0_, then all species *x* _*j*_ strictly causal to *x*_*i*_ are in *I*_0_. Considering the union of the sources also leads to an autonomous pair, because unions of autonomous pairs are autonomous pairs. The minimality follows from the lemma.

In the case when the reachability radius *r* is finite the hierarchical levels (*I*_*m*_, *J*_*m*_), 0 ≤ *m* ≤ *d* − 1 of a r-NHD are not autonomous in the strict sense of Definition 1, but only approximately. The approximation consists in considering that influences fade away with the distance the interaction graph and neglect all influences transmitted along paths longer than *r*.

## 4 The BlockBioNet pipeline

We demonstrate how to combine the concepts introduced in the previous sections to establish a hierarchical block decomposition. The algorithmic workflow, called **BlockBioNet**, consists of several steps listed in the Algorithm 1 and illustrated in the Figure 2.

**Figure 2:**
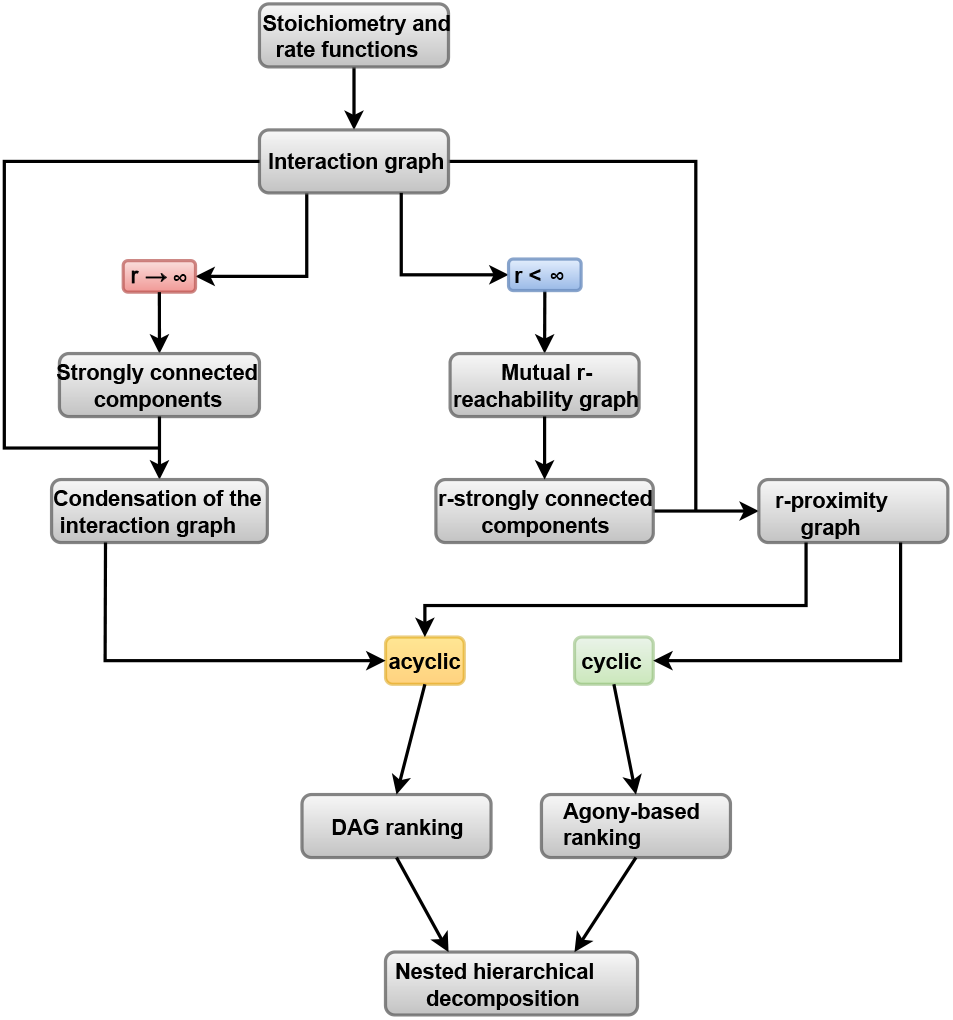
Overview of the BlockBioNet nested hierarchical decomposition algorithm.

### 4.1 Main algorithm

**BlockBioNet**, developed for CRNs, utilizes the interaction graph of these networks. The method is highly versatile and can be used to identify the hierarchical structure of other systems representable as a directed graph, such as social networks [28], transportation networks [29], and ecological networks [30]. Below, we offer a detailed description of the decomposition algorithm.

The first step involves constructing the interaction graph 𝒢. This graph is built using information about reactions and the species involved, including reactants, products, and modifiers, based on Definition 3.

The next steps depend on the chosen value of the reachability radius *r*, which leads to two key cases: *r <* ∞ and *r*→ ∞. The two cases are treated using different algorithms. The case *r*→ ∞ corresponds to strict causality and leads to a-NHDs of autonomous pairs. The case *r <* ∞ leads to r-NHDs.

#### 4.1.1 Finite reachability radius

We first consider the case when the reachability radius is finite, i.e., *r <* ∞. In this case, the next step is to construct the *r*-proximity graph.

If the *r*-proximity graph 𝒬 is acyclic (DAG), we proceed by constructing a r-NHD using hierarchical graph traversal of the proximity graph. The lowest level subset *I*_0_ is the union of the sources of the *r*-proximity graph 𝒬, i.e., the *r*-SCCs with no incoming connections. The corresponding reaction subset 𝒥_0_ comprises all reactions producing or consuming species from *I*_0_. The next level, *I*_1_, is obtained by adding to *I*_0_ all blocks that receive direct connections only from the sources, and so forth. This approach provides hierarchical ranks for all the r-blocks. The species inherit the hierarchical rank of the r-block they belong to. At each level *m*, the reaction subset 𝒥_*m*_ consists of all reactions that have a nonzero net effect on the species in *I*_*m*_.

In the case where the *r*-proximity graph 𝒬 has cycles, we use agony to determine the hierarchical ranks of *r*-SCCs, as described in Section 2.2.

We then establish a r-NHD in exactly the same way as in the previous case.

#### 4.1.2 Infinite reachability radius

When the reachability radius *r* approaches infinity, the *r*-SCCs of the interaction graph 𝒢 reduce to the standard SCCs of 𝒢. In this scenario, the *r*-proximity graph 𝒬 constructed from 𝒢 is equivalent to the condensation of 𝒢, representing the quotient graph of 𝒢 based on its SCCs.

Since, the condensation 𝒬 of the interaction graph 𝒢 is always acyclic, the hierarchical graph traversal leads directly to a r-NHD.

According to the Proposition 1, in this case, the r-NHD is a minimal a-NHD.

These steps collectively define a hierarchical block decomposition for a given CRN. This method organizes the network into blocks of species and their associated reactions, providing a structured approach to analyze and understand the hierarchical organization of complex chemical systems.

### 4.2 Complexity estimates

The **BlockBioNet** pipeline employs two slightly different algorithms for cases where *r* is finite and infinite. When *r* is finite, the block decomposition involves three steps: determining the interaction graph 𝒢, constructing the mutual *r*-reachability graph ℋ, and computing the connected components of ℋ. When *r* is infinite the block decomposition involves only two steps: determining the interaction graph 𝒢, and computing the SCCs of 𝒢.

In order to compute the interaction graph, one needs to process each reaction and determine which species determine its reaction rate and which species are changed by the reaction. Considering that there are *n*_*i*_ reactants, products and modifiers for the *i*th reaction, one has, in the worst case, 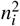 potential interactions resulting from this reaction. Therefore the overall time complexity for computing the interaction graph is *O* ((max_*i*_ *n*_*i*_)^2^*n*), where *n* is the number of reactions.

The *r*-reachability graph ℋ is computed by evaluating the shortest distances between species in the interaction graph 𝒢. This is performed using the Floyd-Warshall algorithm [31–33] whose time complexity is *O*(*s*^3^), where *s* is the number of species.

The connected components of the undirected graph *H* can be computed using Breadth-First Search (BFS), which operates in *O*(*s* + *e*) time, where *e* denotes the number of edges in the graph *H*. In the worst-case scenario, the number of edges *e* can be as large as *s*^2^, leading to a complexity of *O*(*s*^2^). Similarly, the SCCs of the directed graph *G* can be determined using Tarjan’s algorithm, which also runs in *O*(*s* + *e*) time. Here, the worst-case scenario for the number of edges *e* is again *s*^2^, resulting in an overall complexity of *O*(*s*^2^). Therefore, in both cases, the complexity is *O*(*s*^2^).

Thus, the overall time complexity is *O* ((max_*i*_ *n*_*i*_)^2^*n* + *s*^3^) for the finite *r*-reachability case, and *O* (max_*i*_ *n*_*i*_)^2^*n* + *s*^2^ for the infinite *r*-reachability case.

### 4.3 Python implementation

We have developed **BlockBioNet** as a Python package designed for the automated computation of a-NHDs and r-NHDs for CRNs. The package takes as input an SBML file representing the network and a specified reachability radius. It produces the a-NHD, the detailed model information, including the count of species, reactions, edges within the interaction graph, *r*-SCCs, species count per component, and the execution time, that are saved in JavaScript Object Notation (JSON) format within a directory chosen by the user. Furthermore, the package outputs a SBML level 3 description of the model, with blocks added as SBML groups of species and reactions.

Upon execution of the code, a pop-up window initiates, allowing the user to select one or multiple SBML files. Each selected model undergoes identical processing steps. Once an SBML file is chosen, a subsequent window prompts the user to specify the reachability radius. Following input submission, the code systematically progresses through computations to generate the interaction graph, mutual *r*-reachability graph, *r*-SCCs, *r*-proximity graph, agony-based hierarchical ranking, and ultimately identifies the species-reactions pairs that compose the r-NHD.

## 5 Application to biological networks

### 5.1 Decomposing real–life computational models

We demonstrate our decomposition method with a detailed, step-by-step application to real-world computational models, illustrating how the nested hierarchical decomposition is obtained. In particular, we analyze a computational model of the Mitogen-Activated Protein Kinase (MAPK) signaling pathway developed in [34] and simplified in [5] by Michaelis-Menten type reduction, providing a thorough explanation of each step in the decomposition. Two additional examples are included in the supplementary material, where we omit the procedural details and present only the graphical visualizations of the decomposition process.

The reactions involved in the considered computational model of MAPK signalling pathway are detailed below:

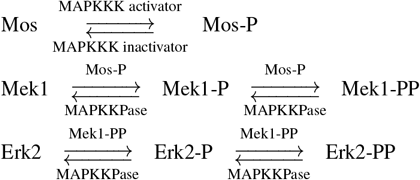

The graphical visualization of the chemical reactions is presented in Figure 3a.

**Figure 3:**
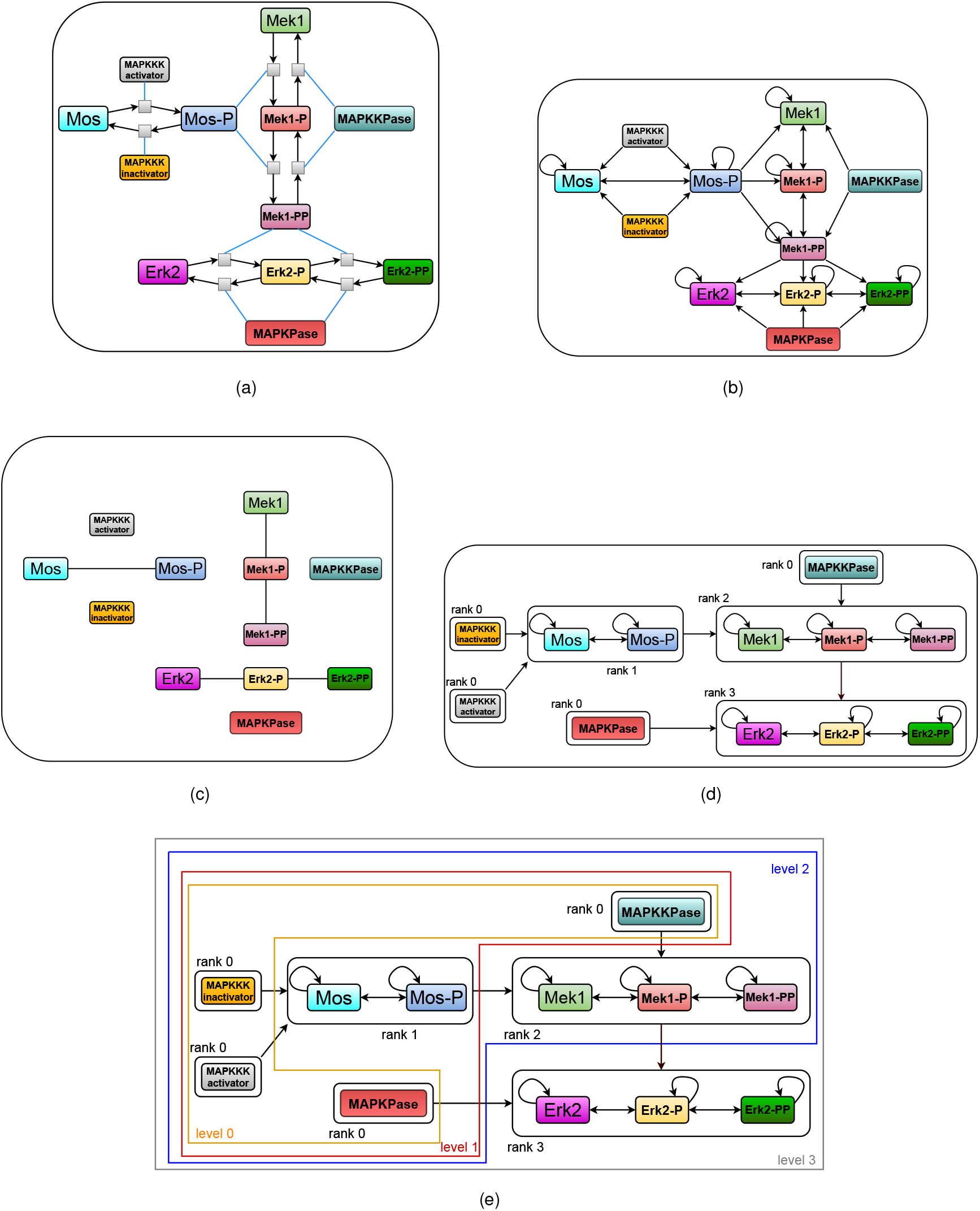
Graphical visualization of the model of the MAPK signaling pathway, and the major steps leading to its nested hierarchical decomposition demonstrated. (a) Graphical visualization of the CRN model as a bipartite oriented graph with two type of nodes: reactions (small squares) and species (rectangles). Black edges indicate that species are consumed (edges originate from species) or produced (edges originate from reactions). Blue edges represent modifier interactions, in which a species influences the rate of a reaction; in this case, the implicit edge orientation from species to reaction is not shown. (b) Interaction graph. (c) Mutual *r*-reachability graph. (d) *r*-proximity graph. (e) r-NHD construction.

According to the first step, we determine the interaction graph 𝒢 of the CRN. The set of vertices corresponds to the species involved. Since the CRN includes ten species, the interaction graph has ten vertices, each representing one species. We construct the edges of the interaction graph 𝒢 as follows.

For example, consider the first reaction: it consumes a molecule of Mos and produces a molecule of Mos–P, with a reaction rate that depends on Mos and the MAPKKK activator. Here, Mos acts as the substrate and the MAPKKK activator as a modifier. Consequently, in the interaction graph 𝒢, there are directed edges from both Mos and the MAPKKK activator to Mos–P. Following this approach, we can construct all other edges of the interaction graph 𝒢, as represented in Figure 3b.

We selected a reachability radius of one, noting that a larger radius would result in blocks containing more species. Therefore, the second step in our decomposition procedure is the determination of the mutual *r*-reachability graph. The mutual *r*-reachability graph is constructed as follows: there is a directed edge from Mos to Mos-P in the interaction graph, and similarly a directed edge from Mos-P to Mos, this implies 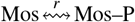. Consequently, in the mutual *r*-reachability graph, there is an undirected edge between Mos and Moss–P. Through analogous reasoning, we complete the mutual *r*-reachability graph, which is depicted in Figure 3c. It is important to note that some species are isolated in this graph, as they are not mutually *r*-causal to any other species. The *r*-SCCs of the interaction graph are the connected components of the mutual *r*-reachability graph, i.e. {MAPKKK activator}, {MAPKKK inactivator}, {MAPKPase}, {MAPKKPase}, {Mos, Mos–P}, {Mek1, Mek1–P, Mek1–PP}, {Erk2, Erk2-P, Erk2–P }.

The next step in our procedure is to determine the *r*-proximity graph, which is the quotient graph defined by the set of *r*-SCCs. In this graph, each node corresponds to an *r*-SCC. Since there is a directed edge from species Mos–P to Mek1 in the interaction graph and these species are in different *r*-SCCs, the *r*-proximity graph contains a directed edge from the node corresponding to the *r*-SCC containing Mos–P to the node corresponding to the *r*-SCC containing Mek1. We construct all the other edges in a similar manner, resulting in the final *r*-proximity graph, which is graphically represented in Figure 3d. We observe that the *r*-proximity graph is acyclic. Consequently, we apply the agony-based node ranking method established in [19] to rank the nodes in the *r*-proximity graph. The ranks are also included in Figure 3d.

We establish a r-NHD from this ranking as follows: The lowest level *I*_0_ comprises the lumping of *r*-SCCs with zero rank, namely

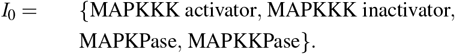

The reaction set at this level is

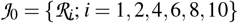

as these reactions involve species in *I*_0_.

The subsequent level, *I*_1_, extends *I*_0_ to include *r*-SCCs with rank one. Here, there is only one such a *r*-SCC which is {Mos, Mos–P}, Thus:

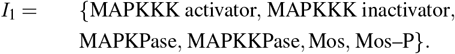

The corresponding reaction set at the second level is

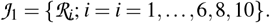

The next level, *I*_2_, includes *I*_1_ extended by *r*-SCCs with rank two. The singular such *r*-SCC is {Mek1, Mek1–P, Mek1–PP}, thus, *I*_2_ is given by

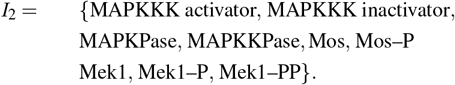

The corresponding reaction set

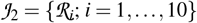

encompasses reactions influenced by species in the third hierarchical level.

Finally, the last level, *I*_3_, includes *I*_2_ extended by *r*-SCCs with rank three. The only such *r*-SCC is {Erk2, Erk2–P, Erk2–PP }, thus, *I*_3_ is given by

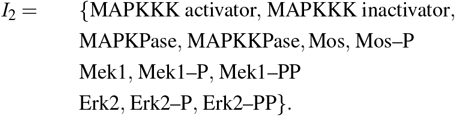

Naturally, the corresponding reaction set is 𝒥_3_ = 𝒥_2_. The aggregation of *r*-SCCs into hierarchical levels is illustrated in Figure 4. Applying this method, we systematically determined network hierarchies, shown in Figure 3e with colored boxes for clarity.

**Figure 4:**
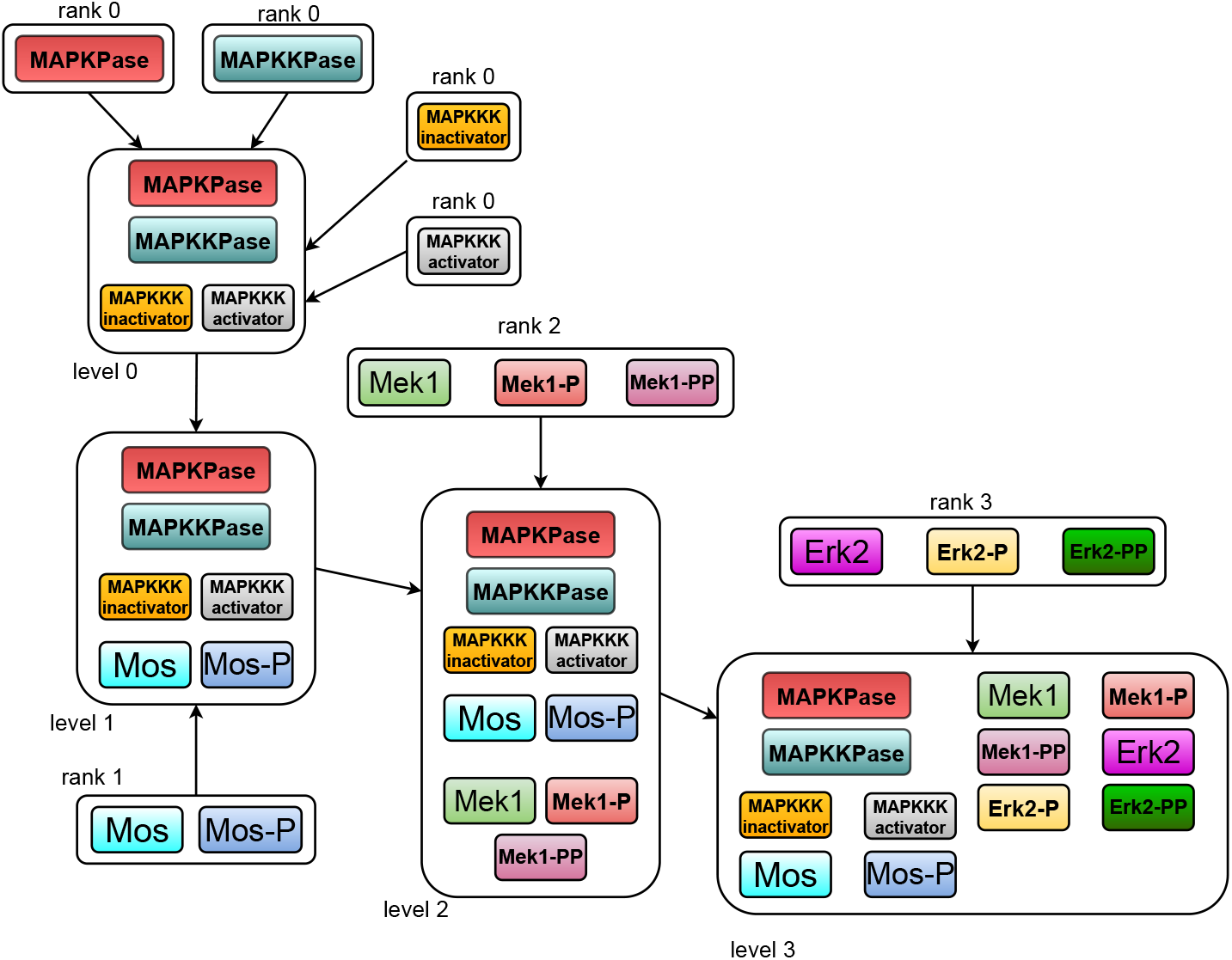
Graphical visualization of component lumping for the model of the MAPK signaling pathway

### 5.2 Benchmarking using BioModels database

We applied our hierarchical decomposition method to 1057 CRN models extracted from the **Biomodels** database [35]. In order to study the effect of changing *r* on the decomposition we have performed the decomposition for values 1 ≤ *r*≤ 5.

As shown in Figure 5a, the algorithm has an average complexity *O*(*s*^1.98^), or equivalently *O*(*e*^4*/*3^) where *s, e* are the numbers of species, and edges in the interaction graph, respectively. This is true for all values of *r* considered (the data shown in Figure 5a, gathers values 1 ≤ *r* ≤ 5). This result is better than the theoretical limits. The excellent performance can be explained on one hand by the fact that CRN models used in systems biology have sparse interactions. On the other hand, unlike typical implementations of hierarchical decompositions, we do not solve the computationally expensive agony optimization directly on the interaction graph, but on the *r*-proximity graph or the condensation of the interaction graph, which is less complex.

**Figure 5:**
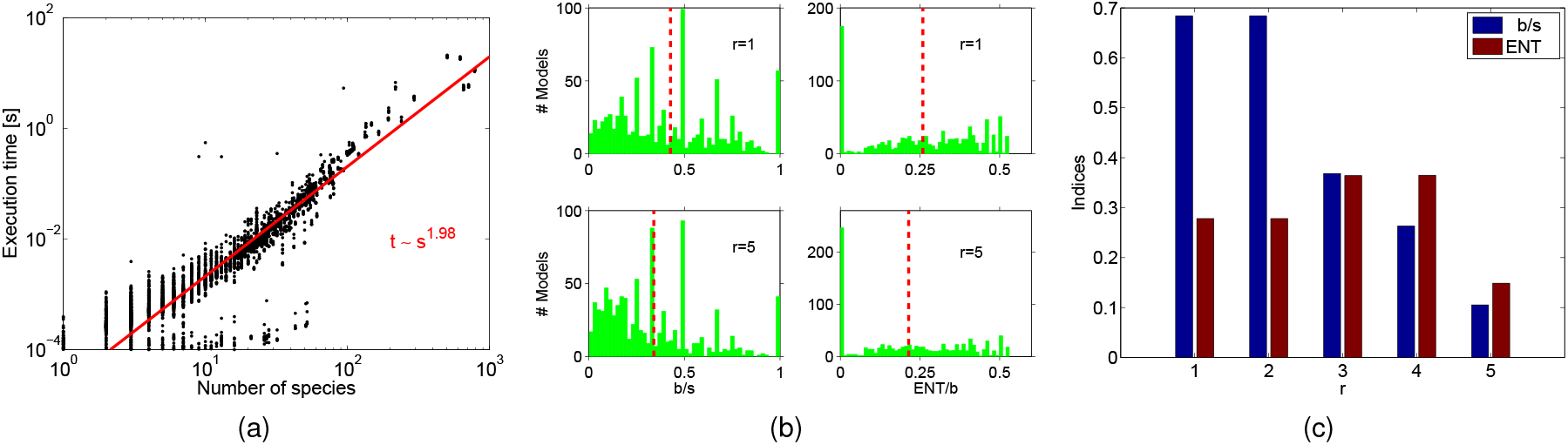
Benchmarking **BlockBioNet** on the **Biomodels** database [35], using a total of 1057 models. (a) Execution time vs interaction graph edge count; we gather results for 1≤ *r* ≤ 5. (b) Distributions of number of blocks and specific entropy; mean values are indicated by the red dotted lines. (c) *r* selection procedure for the model BIOMD0000000096; the selected value *r* = 3 maximizes the number of blocks at maximum specific entropy.

In order to asses the quality of the decomposition we defined two indices. The first one is the inverse block size, defined as number of blocks *b* divided by the number of species *s*:

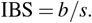

IBS is a rational number between 0 and 1 that represents the inverse mean number of species per block.

The second index is the specific entropy (entropy per block) of the block decomposition

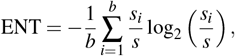

where *s*_*i*_ is the number of species in the block *i*.

A large value of ENT indicates that the species are relatively uniformly distributed across different blocks, whereas a small value indicates non-uniform distributions.

The values of the two indices for different models are shown in Figure 5b. We can notice that larger values of *r* corresponds to smaller means of the two indices, suggesting that increasing *r* tends to decrease both the number of blocks and the specific entropy.

IBS is monotonously decreasing with *r* for all models. However, ENT is either decreasing with *r* or has a maximum for one or several values of *r*, as shown in Figure 5c. This type of behavior suggests a method to select the optimal value of *r*. For applications such as hierarchical optimization [5], a good decomposition has a large entropy (species uniformly distributed in blocks). For the same entropy, having more blocks is better because these blocks are smaller and therefore easier to analyze. Thus, the selection criterion for *r* is as follows:

- find those *r* that maximize the specific entropy ENT.
- choose among them the one corresponding to the largest IBS.

### 5.3 Block Visualization using CRNPlot

Visualization of biochemical networks is an important topic in bioinformatics, helping in the understanding of the intrinsic complexity of biochemical data. Many databases, such as KEGG, provide static biochemical pathway maps as bitmaps. However, tools for dynamic visualization, enabling automated graphical representations of pathways based on database queries or automated model generation, have yet to be developed.

There have been numerous attempts to develop pathway visualization tools. [36] lists 51 network visualization resources, most of which are either general-purpose or specialized for transcriptional and protein-protein interaction graphs. A few resources have been developed for metabolic networks, that are CRNs, but these are typically either static, relying on predefined maps or manually adjustable layouts, or not yet implemented into functional tools. For instance, the algorithmic approach proposed by [37], which decomposes metabolic pathways into linear, tree-like, and cyclic components and arranges them using spring embedding layouts, is not available as a tool. Moreover, while this method was tailored for metabolic networks, it is not broadly applicable to all CRNs. Some visualization tools for protein-protein and transcriptional interactions incorporate blocks and hierarchical levels [38,39]. However, these tools rely on community detection algorithms and do not extend to species-reaction graphs.

In this paper we have proposed a theoretical framework for multiple graphical representations of CRNs: species-reactions graphs, interaction graphs, hierarchical graphs. We developed **CRNPlot** that uses block decomposition of CRNs, generated by **BlockBioNet** as a SBML level 3 file, to visualize the bipartite species-reaction graph, the interaction graph and the r-proximity graph. **CRNPlot** uses Graphviz for automated generation of directed graphs layouts, represented in dot language. In a first step, a layout is generated for the boxes containing the blocks. Then, the boxes are filled with the block species and reactions.

An illustration of **CRNPlot** application is shown in the Figure 6. For a model of mTOR/AMPK signaling pathway crosstalk [40] we have generated graphs for different values of the scale parameter *r*. Contrary to hierarchical optimization applications, where a large number of blocks (corresponding to a small *r*) is preferred, larger and fewer blocks may be a better choice for visualization (corresponding to larger *r* values), even if the block sizes are not uniform (i.e., ENT is low). In this case, proposing objective criteria is challenging; therefore, the choice is left to the user.

**Figure 6:**
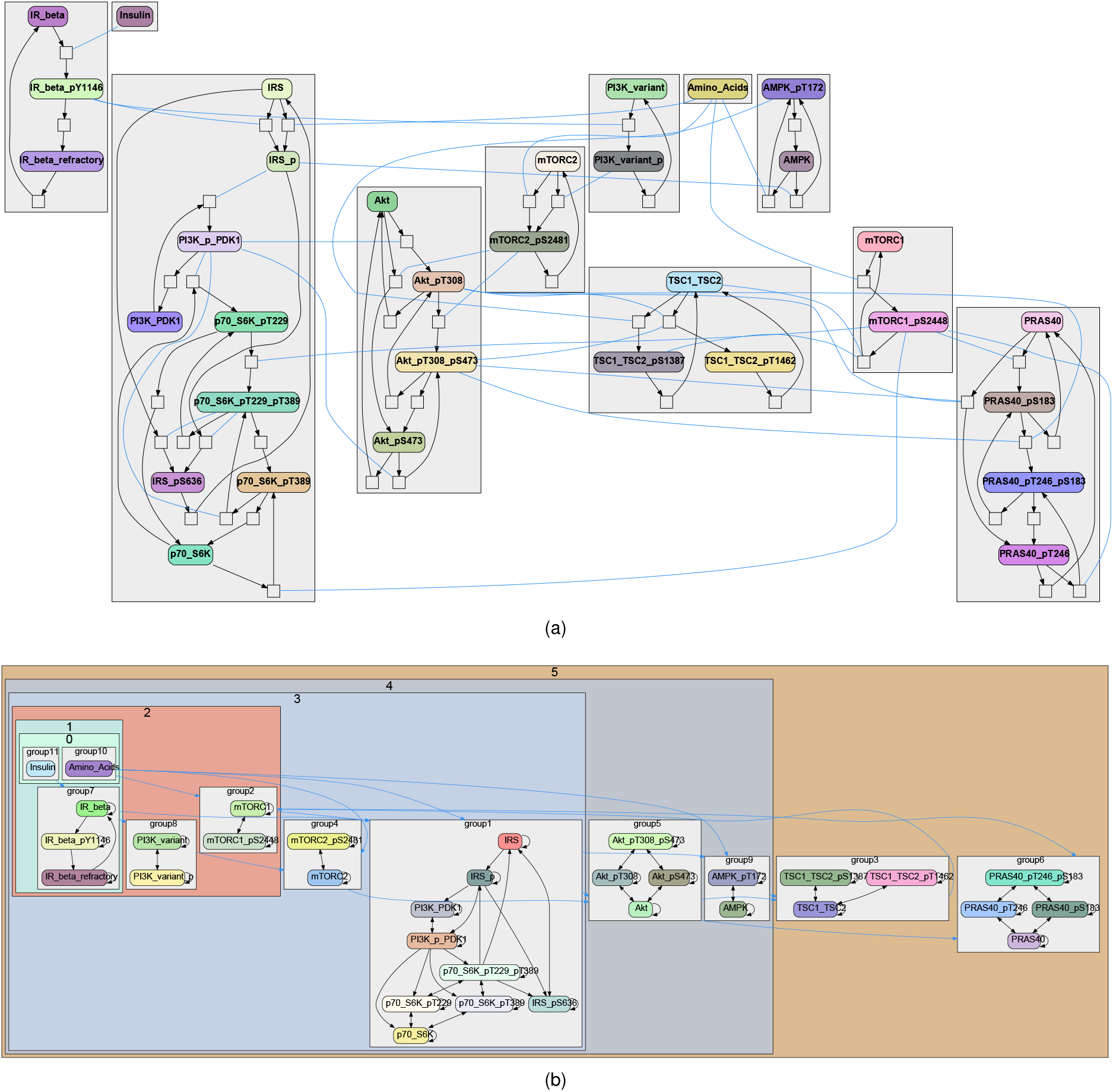
Block decomposition generated by **BlockBioNet** and visualized by **CRNPlot** for a mTOR/AMPK signaling pathway crosstalk model [40]. The model is available in the **Biomodels** database [35] (identifier BIOMD0000000640). a) species-reaction graph with boxes defining *r*-SCCs; a blue edge connecting a species and a reaction indicates that the species acts as a modifier of the reaction. b) r-NHD construction. The reachability radius is *r* = 2.

The block decomposition of the model emphasizes the interactions between the model components. Figure 6b) shows the multiple inputs of the amino acids on the mTOR network, one of the original’s work main results.

## 6 DISCUSSION

High-throughput experimental methods [1–3, 41] and automated model building [42–44] yield large reaction networks lacking block organization. Similar graphs are already common in transcriptional control networks. These dense networks contrast with the much more modular diagrams typical of signaling pathways. The current study provides a theoretical and practical basis for deriving such modularity. In the traditional approach to signaling pathways, genetics was used to define key molecules implicated in a given function, and targeted biochemistry helped to fill out the reactions [45]. Thus signaling pathways were functionally defined. **BlockBioNet** is remarkably effective in reconstructing these pathway blocks based on topology rather than functional information. Thus a first key step in biological insight is obtained by grouping reactions into pathways, each of which has an implicit functional role. A second important insight is the topology of information flow. Having a more coarse-grained, block-level description of a dense reaction network assists in tracing signal propagation, and identifying motifs such as feedback loops. Numerous motif-finders are present in the literature, such as **ReactionFlow** [46], **BiNoM** [47], or **MORO** [48]. In comparison to these, and when combined with the network block visualization tool **CRNPlot, BlockBioNet** provides insight in two ways: it performs coarse-graining and explicitly reports loops.

More generaly, our methods can be used to study causality in general networks. Causality studies can be used not only to understand model functioning but also for machine learning applications. In particular, as we recently proposed [5], hierarchical optimization, which involves decomposing large mechanistic computational models into blocks and optimizing them progressively, starting with input blocks and following causality paths, significantly reduces the computational burden, as each step involves solving a simpler problem.

Breaking problems into smaller parts is a general approach to overcoming the complexity barrier of other hard problems in computational biology such as network alignement, motif discovery, combinatorial optimization problems (minimal network hit sets and cuts). Decomposition techniques were used for heuristic approaches and approximated solutions for several NP-complete problems such Boolean Satisfiability (SAT) [49, 50], SAT modulo theorys (SMTs) [51], integer and mixed integer programming [52].

Our approach has certain limitations, which we plan to address in future work.

In this approach, all interactions are assumed to have equal importance, which does not always reflect real-world conditions. This limitation can be addressed by assigning weights to the edges of the interaction graph. Consequently, a natural extension of our method would be to generalize the computation of agony so that edges violating the hierarchy are penalized in proportion to their weights.

We can also observe that, depending on the chosen reachability radius *r*, the resulting groups may be unbalanced, as some subsets of species interact more densely than others. A pathological case arises when each species lies within a distance *r* of every other species, resulting in a single block encompassing all species. Therefore, it may be useful to impose constraints on the decomposition, for example, by limiting the maximum size or the number of groups, to ensure that the resulting subproblems remain tractable.

Finally, networks with cycles inevitably face a circular dependency problem, as signaling flow is not well-defined and can proceed in both directions. In such cases, the block decomposition (the r-proximity graph) is biologically informative as it provides coarse-grained representation of the network, emphasizing blocks, interaction between blocks and feedback. The nested hierarchical structure may be artificial, with lowest ranks arbitrarily chosen in cycles (see Supplementary Figures), but it is still useful for computational applications such as hierarchical optimization [5].

## Acronyms

SCC: *r*-Strongly Connected Component
SBML: Systems Biology Markup Language
JSON: JavaScript Object Notation
CRN: Chemical Reaction Network
ODE: Ordinary Differential Equation
a-NHD: Autonomous Nested Hierarchical Decomposition
r-NHD: *r*-Causal Nested Hierarchical Decomposition
SAT: Satisfiability
SCC: Strongly Connected Component
SMT: Satisfiability Modulo Theory

## Code Availability

The Python package developed for this work is available at the following links: https://github.com/tambysatya/crnplot, https://github.com/manvelgasparyan/BlockBioNet.

## Acknowledgements

This work has been supported by CEFIPRA grant 68T08-3.

## Supplementary Figures

**Figure S1:**
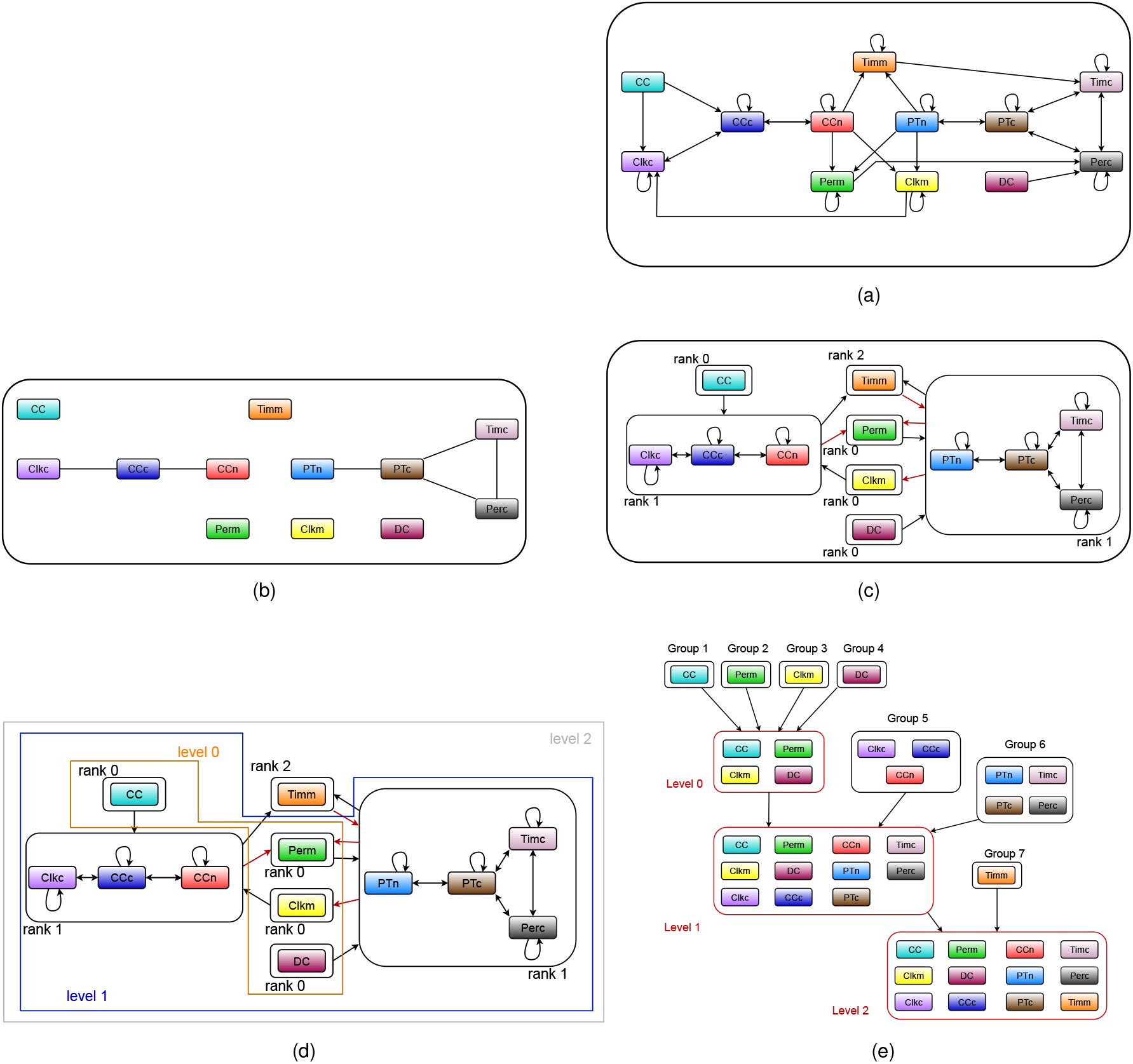
Hierarchical block decomposition of the model Ueda2001 Circadian Clock model (BioModels ID: BIOMD0000000022), as described in [53]. The model described the molecular feedback loops underlying circadian rhythms in Drosophila. It captures how interactions between core clock proteins generate self-sustained oscillations with ~ 24-hour periodicity. (a) Graphical visualization of the model. (b) Interaction graph. (c) Mutual *r*-reachability graph. (d) *r*-proximity graph. (e) Component lumping. (f) r-NHD construction. The edges penalized by agony are drawn in red. The r-proximity graph contains loops, one of which includes the variable PERm, representing the mRNA of the *per* gene, and PTn, representing the corresponding protein. Facing the circular dependency between these two variables, the agony algorithm randomly assigned PERm and PTn as rank 0 and rank 1, respectively. However, choosing the opposite ranking would be equivalent.

**Figure S2:**
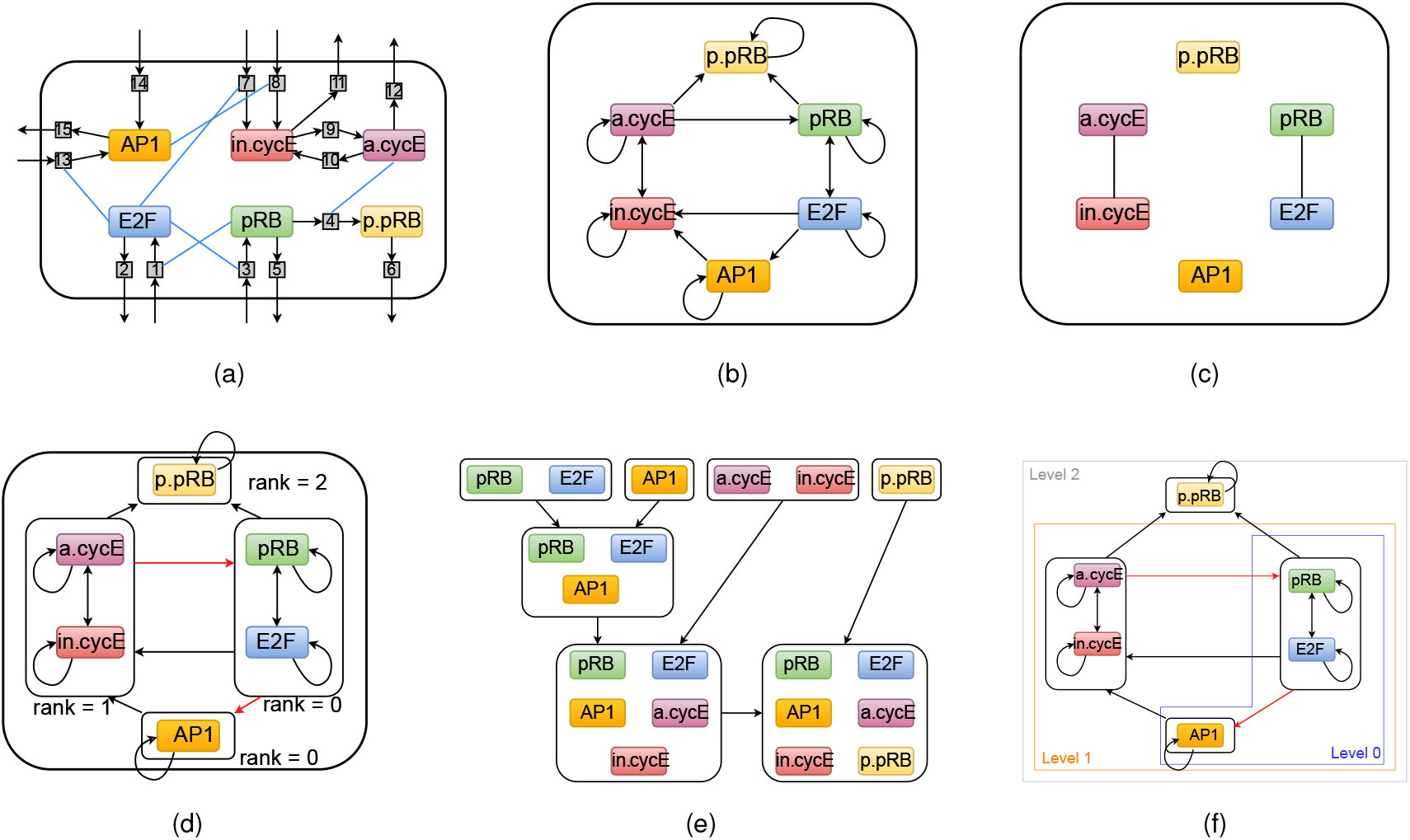
Model of the G1 phase and the G1/S transition in the mammalian cell cycle, as described in [54]. (a) Graphical visualization of the model. (b) Interaction graph. (c) Mutual *r*-reachability graph. (d) *r*-proximity graph. (e) Component lumping. (f) r-NHD construction. The edges penalized by agony are drawn in red. The r-proximity graph contains many loops. Although the blocks are informative, the levels may seem artificial.

